# Deep Learning-based Optical Aberration Estimation Enables Offline Digital Adaptive Optics and Super-resolution Imaging

**DOI:** 10.1101/2023.10.27.564396

**Authors:** Chang Qiao, Haoyu Chen, Run Wang, Tao Jiang, Yuwang Wang, Dong Li

## Abstract

Optical aberrations degrade the performance of fluorescence microscopy. Conventional adaptive optics (AO) leverages specific devices, such as the Shack-Hartmann wavefront sensor and deformable mirror, to measure and correct optical aberrations. However, conventional AO requires either additional hardware or a more complicated imaging procedure, resulting in higher cost or a lower acquisition speed. In this study, we proposed a novel space-frequency encoding network (SFE-Net) that can directly estimate the aberrated point spread functions (PSFs) from biological images, enabling fast optical aberration estimation with high accuracy without engaging extra optics and image acquisition. We showed that with the estimated PSFs, the optical aberration can be computationally removed by deconvolution algorithm. Furthermore, to fully exploit the benefits of SFE-Net, we incorporated the estimated PSF with neural network architecture design to devise an aberration-aware deep-learning super-resolution (DLSR) model, dubbed SFT-DFCAN. We demonstrated that the combination of SFE-Net and SFT-DFCAN enables instant digital AO and optical aberration-aware super-resolution reconstruction for live-cell imaging.

## 1. INTRODUCTION

Fluorescence microscopy has been widely used as a powerful tool for visualizing various biological structures and bioprocesses in fixed or live specimens. Ideal optical imaging relies upon the high-quality focusing of excitation light and accurate detection of the emission light from the fluorescent sample. However, both the optics in the microscope and the biological samples being investigated can introduce aberrations, thus causing degradation in resolution, loss of fluorescent photons, and deterioration of signal-to-background-ratio (SBR), etc. For example, the optics manufacture deficiency or misalignment of optical elements in the imaging system may cause certain aberrations such as spherical and coma aberration, and the refractive index inhomogeneities of biological specimens will bring about more complicated aberrations. Moreover, microscopes with high numerical apertures (NA), especially the super-resolution microscopy, are more sensitive to aberrations, because the high-NA objectives are more susceptible to high-order aberrations [1]. To detect and correct these optical aberrations, a large number of adaptive optics (AO) technologies have been explored in the last two decades [2].

The implementation of AO generally involves two main components: aberration detection and aberration correction. To measure optics- or sample-induced aberrations, both direct and indirect wavefront sensing methods were developed [1-3]. Direct wavefront sensing methods utilize a dedicated wavefront sensor, mostly the Shack-Hartmann sensor, along with an additional light path for aberration detection. In contrast, the indirect wavefront sensing methods characterize aberrations without specific wavefront sensors but determine them computationally from repetitive acquisitions with either zonal or modal approaches [2]. In recent years, deep neural networks are applied to directly estimate aberrations from the optical images of point sources [4-6]. However, these methods are limited to the scenarios where there is the guiding star or single-molecule emitters in the biological samples. Once the aberrations are known, wavefront corrective devices, mostly the spatial light modulators (SLMs) and deformable mirrors (DMs), are utilized to compensate for the measured aberrations by reshaping the wavefronts [1-3]. In consequence, conventional AO methods have to rely on additional optical device or iterative acquisitions to measure and then eliminate the optical aberration, which complicates the optics, imaging procedures, and computation. To overcome these limitations, the development of digital adaptive optics (DAO) has allowed for the computational detection and correction of optical aberrations for light-field microscopy (LFM) [7, 8] in an offline manner, which, however, is only applicable for the certain imaging modality, i.e., LFM.

In optical imaging systems, the image quality as well as the aberrations is typically characterized by their point spread functions (PSF), which is implicitly encoded in any specimen patch of the microscopic image. Inspired by the understanding, we devised a space-frequency encoding network (SFE-Net), which is trained to directly extract the PSF with aberrations from a single microscope image. Our results show that the proposed SFE-Net is able to estimate optical aberrations composed of up to 18 Zernike polynomials with high accuracy directly from images of various biological specimens, and the corresponding aberrations can be substantially eliminated via the deconvolution algorithm resorting to the estimated PSF. To further enhance the resolution while removing the optical aberrations for biological images, we integrated the PSF priors into the deep-learning super-resolution (DLSR) neural network architecture design, and devised the spatial feature transform-guided (SFT) deep Fourier channel attention network (SFT-DFCAN). We showed that by leveraging PSF information estimated from SFE-Net, the SFT-DFCAN can be trained to digitally eliminate the aberrations and super-resolve the fine structures of specimens directly from the aberrated images, which substantially outperforms its backbone DFCAN architecture [9]. Finally, we demonstrated that the SFE-Net and SFT-DFCAN enable fast, accurate aberration estimation and correction, as well as computational super-resolution, in long-term live-cell imaging experiments.

## METHODS

### A. Training Data Generation

The training data for SFE-Net, SFT-DFCAN, and other deep-learning models compared in this study was generated in a semi-synthetic manner using our previously published dataset BioSR [9]. Specifically, we utilized the ground truth structured illumination microscopy (GT-SIM) images from BioSR as the biological fluorescence specimens. These images were intentionally degraded according to the optical imaging model, which can be expressed as follows:

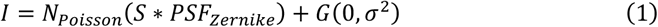

where *I* represents the aberrated wide-field (WF) image captured by the optical imaging system, *S* denotes the biological specimens, i.e., the GT-SIM images, ∗ signifies the convolution operator, *N*_*Poisson*_(·) represents the Poisson recorruption, *G*(0, *σ*^2^) denotes the Gaussian white noise with a mean of zero and a variance of *σ*^2^, and *PSF*_*Zernike*_ refers to the aberrated points spread functions, whose pupil functions are constructed by Zernike polynomials 4-18 (Wyant ordering) with random amplitudes ranging from −1.0 to 1.0 for each order. These functions can be mathematically formulated as follow:

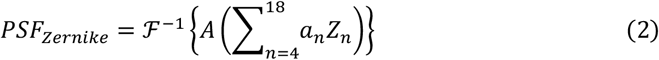

where *Z*_*n*_ and *a*_*n*_ represent Zernike polynomials and amplitudes of order *n*, respectively. *A*(·) denotes a circular apodization function, where the radius is determined by the emission wavelength and detection numerical aperture (NA). ℱ^-1^{·} denotes the inverse fast Fourier transformation operator.

During the training procedure, the aberrated PSF images *PSF*_*Zernike*_ were used as targets for PSF estimation network models such as SFE-Net, and the GT-SIM images *S* are used as targets for single image super-resolution (SISR) network models such as SFT-DFCAN.

### B. Network Architecture

The architecture of SFE-Net, as shown in Fig. 1a, consists of a dual-branch encoder and a U-net-based decoder. The encoder network is constituted with two parallel branches: the spatial branch (SB) and frequential branch (FB), which extract deep features in spatial and frequential domain, respectively. In both branches, a modified residual channel attention network (RCAN) [10] with 4 residual groups x 4 residual channel attention blocks is employed as their backbone network architecture (Fig. 1b). In contrast with the SB, the FB begins with a fast Fourier transform layer followed by a modulus operator and a logarithm operator in sequence, so as to encode the image feature into Fourier domain. The output feature maps of SB and FB are concatenated along the channel and then fed into the U-net-based decoder.

**Fig. 1.**
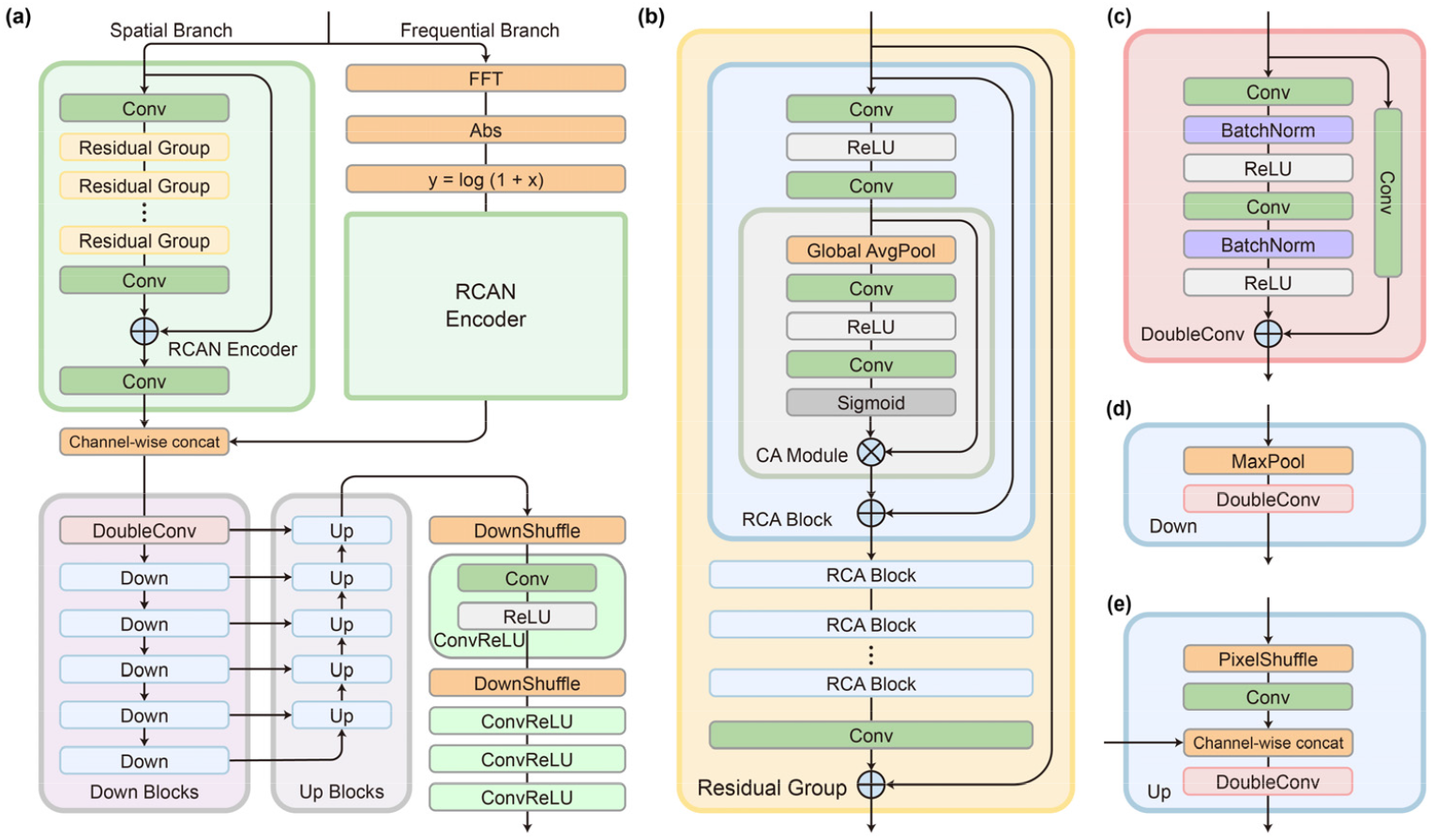
Network Architecture of Spatial-frequency Encoding Network. (a-e) Network architecture of the SFE-Net (a), residual group (b), double convolutional block (c), downscale block (d), and upscale block (e).

The decoder mainly consists of two parts: a U-net feature extractor and a downscale module. We adopt a relatively deep U-net model [11] that begins with a double convolutional block (Fig. 1c) followed by 5 downscale blocks (Fig. 1d) and 5 upscale blocks (Fig. 1e) with 5 skip connections bridging the features of the same scale. In each downscale block, a max pooling layer and a double convolutional block are employed to downscale and extract features. The upscale blocks use the pixel shuffle layer to upscale the feature channels. The output of the U-net is then passed to the downscale module, which consists of two 2× down shuffle layers and four Conv-ReLU blocks. In each Conv-ReLU block, the stride parameter of convolutional layers is set to 2, enabling the downscale module to transform the input feature maps from 132×132 pixels into PSF images of 33×33 pixels.

The overall architecture of SFT-DFCAN is depicted in Fig. 2a, which is modified from our previously proposed state-of-the-art DLSR model DFCAN [9], which is trained to directly transform a WF image to its SR counterpart. Here, to deliver PSF and aberration information to the image SR processing, inspired by the SFTMD model [12], we updated the original Fourier channel attention block (FCAB) into the spatial feature transform-guided FCAB (SFT-FCAB), which could leverage the embedded PSF information to adaptively rescale the spatial features in SFT-RCAB. Specifically, we employed the principal component analysis (PCA) to project the PSF onto a linear space of dimension b. This projected PSF is then stretched into a PSF embedding of size bxHxW, which serves as the input for every SFT-FCAB. In each SFT-FCAB, the PSF embedding are combined with spatial feature maps of the biological structure through two convolution-activation blocks to scale and shift the input feature maps, and subsequently a FCA layer (Fig. 2c) is implemented to perform deep feature extraction and aggregation. Finally, the reconstructed SR image is generated by an up-sampling module, which sequentially consists of a convolution-GELU block[13], a pixel shuffle layer[14], and a final convolutional layer.

**Fig. 2.**
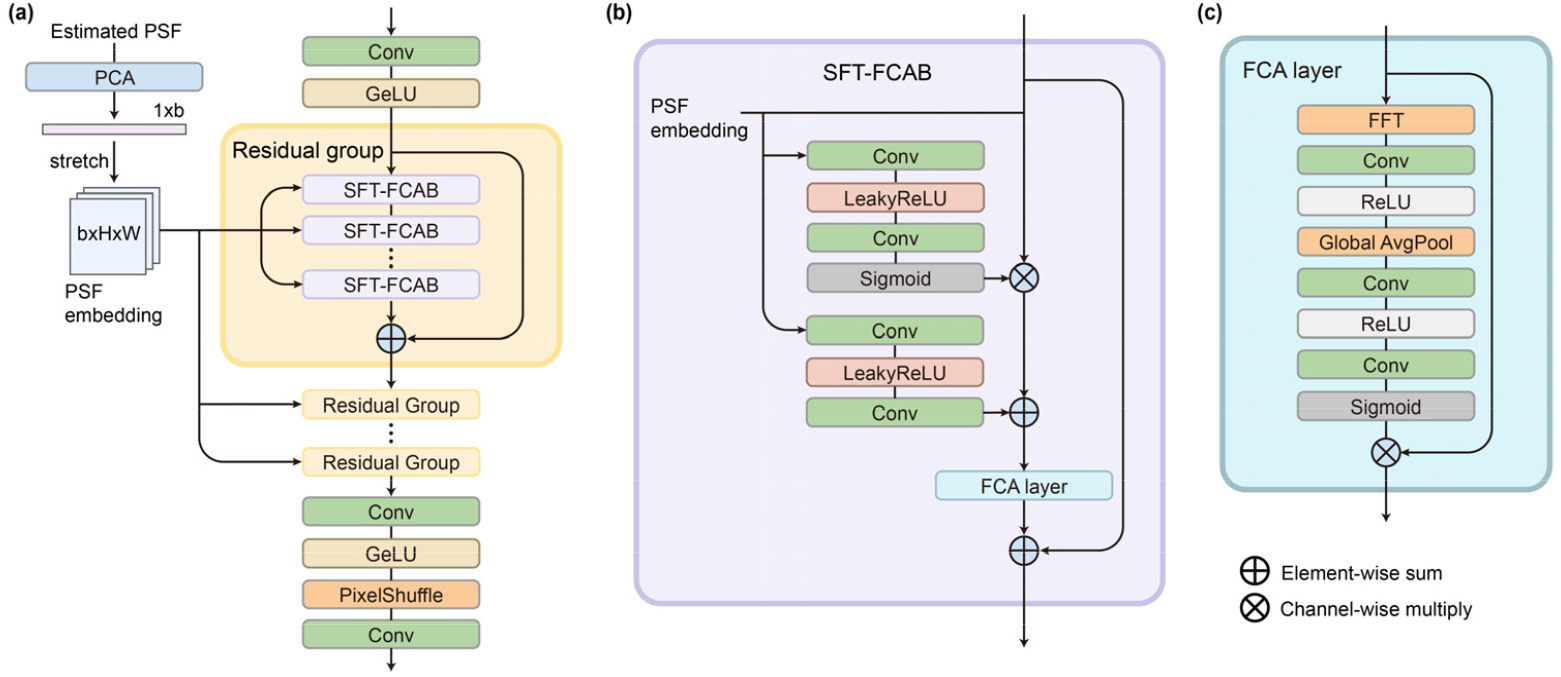
Network Architecture of Spatial Feature Transform-guided Deep Fourier Channel Attention Network (SFT-DFCAN). (a-c) Network architecture of the SFT-DFCAN (a), spatial feature transform-guided Fourier channel attention block (FCAB) (b), and the Fourier channel attention (FCA) layer (c).

### C. Network Training and Data Processing

For training of SFE-Net, we randomly generated pairs of aberrated WF images (132×132 pixels) and theirs corresponding PSF (33×33 pixels) during each iteration following Eq. (1) and Eq. (2) with ∼200 original GT-SIM images of multiple biological specimens, including the hollow clathrin-coated pits (CCPs), the endoplasmic reticulum (ER), and the crisscrossing microtubules (MTs), so as to endow a well generalization capability of the trained model. The overall data augmentation workflow and the training process of SFE-net are shown in Fig. 3. For SISR models such as SFT-DFCAN, we randomly generated triplets of aberrated WF images (132×132 pixels), ground truth PSFs (33×33 pixels), and corresponding GT-SIM images (264×264 pixels) during each iteration as the training dataset. The objective function of both SFE-Net and SISR models is defined as the mean square error (MSE), which quantifies the difference between the network outputs and target images.

**Fig. 3.**
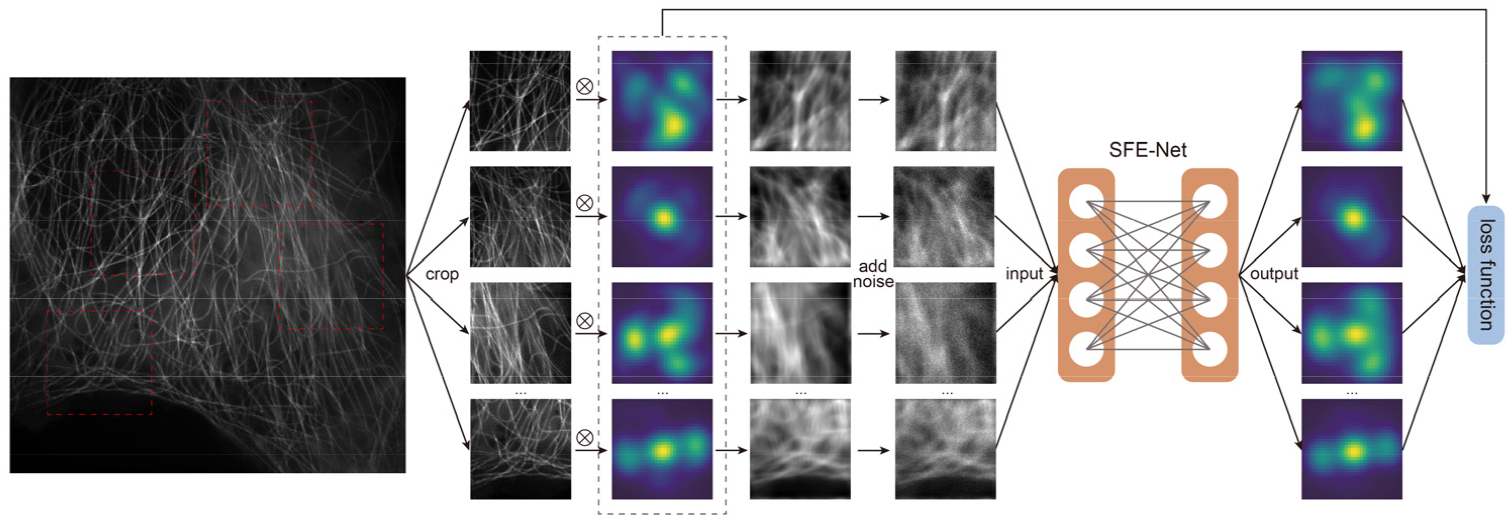
Schematic of the data augmentation and training process of SFE-Net.

The training and inference were performed on a computer workstation equipped with a Interl Xeon(R) Gold 6134 CPU at 3.20 GHz and a NVIDIA RTX 3090 graphic processing card with python v.3.6 and Pytorch 1.12. During the training process, we used the Adam optimizer with an initial learning of 5 x 10^-5^. The learning rate for SFT-Net was decayed by a factor of 0.5 after every ∼10,000 minibatch iterations, while the learning rate of SFT-DFCAN followed a cosine annealing schedule, restarting at every 12,500 minibatch iterations. We adopted a batch size of 4 and 8 for SFE-Net and SFT-DFCAN, respectively. Typically, the total training iterations of SFE-Net and SFT-DFCAN are 150,000 and 500,000, which takes about 16 hours and 30 hours with the RTX 3090 GPU, respectively. In the inference phase, SFE-Net typically takes less than 1.5s (30ms for a single image patch) to generate a PSF matrix (7×7×33×33) by segmenting the input image (512×512) into several patches to capture the spatial variation of optical aberrations. By taking the WF image and estimated PSF as inputs, a well-trained SFT-DFCAN model could reconstruct an aberration-free SR image of 1024×1024 pixels within one second.

## 3. RESULTS

### A. Optical Aberration Estimation via SFE-Net

The PSF encodes substantial and intrinsic information, encompassing optical aberrations and resolution, for both natural images and microscopic images. In recent years, several methods have been developed to estimate the blur kernel of image capture process for natural images [12, 15-17]. However, there have been limited advancements in blind estimation techniques for optical image PSF. The reasons are twofold. Firstly, due to the elaborate optical system and sample scattering, the optical aberration encountered in biological imaging are dramatically heavier than those in commercial camera-based photography. Secondly, estimating the aberrated PSF directly from biological images is essentially an ill-posed problem, rendering it infeasible in intuition. Nonetheless, the image-based estimation of PSF and optical aberration holds substantial benefits for biological imaging, which eliminates the need for a wavefront sensor in AO system, while facilitating digital aberration correction and aberration-aware image super-resolution reconstruction.

In order to address the issues above, we started with exploring several representative supervised or unsupervised kernel estimation algorithm to estimate the kernel, i.e., the aberrated PSF, from biological images. The algorithms included the unsupervised kernel generative adversarial network (KernelGAN) [15, 18], iterative kernel correction (IKC) [12], and supervised mutual affine network (MANet) for spatially variant kernel estimation [17]. To evaluate the performance of these methods, we generated four semi-simulated datasets with aberrated PSF following the procedure outlined in Section A, each constituted by different orders of Zernike polynomials (4-6, 4-8, 4-13, and 4-16). The increasing order range reflects the increasing severity of the ill-posedness in the PSF estimation task. The generated datasets were then utilized to evaluate the performance of existing kernel estimation methods. The results, depicted in Fig. 4a, revealed that the KernelGAN method only generated narrowed anisotropic kernels that significantly deviated from the GT PSF in terms of shape and size. This discrepancy may arise from the multiple kernel constraints in the algorithm, despite we have made great efforts to optimize the weighting scalar of each regularization term. In contrast to KernelGAN, both the IKC and MANet methods consistently produce Gaussian-shaped kernels, regardless of the complexity of training dataset or biological structures. This indicates that these two methods fail to resolve the optical aberrations from the corresponding WF images. In particular, even when we modified the MANet to focus on estimating a spatially consistent kernel, it still could not generate the correct PSF with notable aberrations, possibly due to its relatively simple network architecture.

**Fig. 4.**
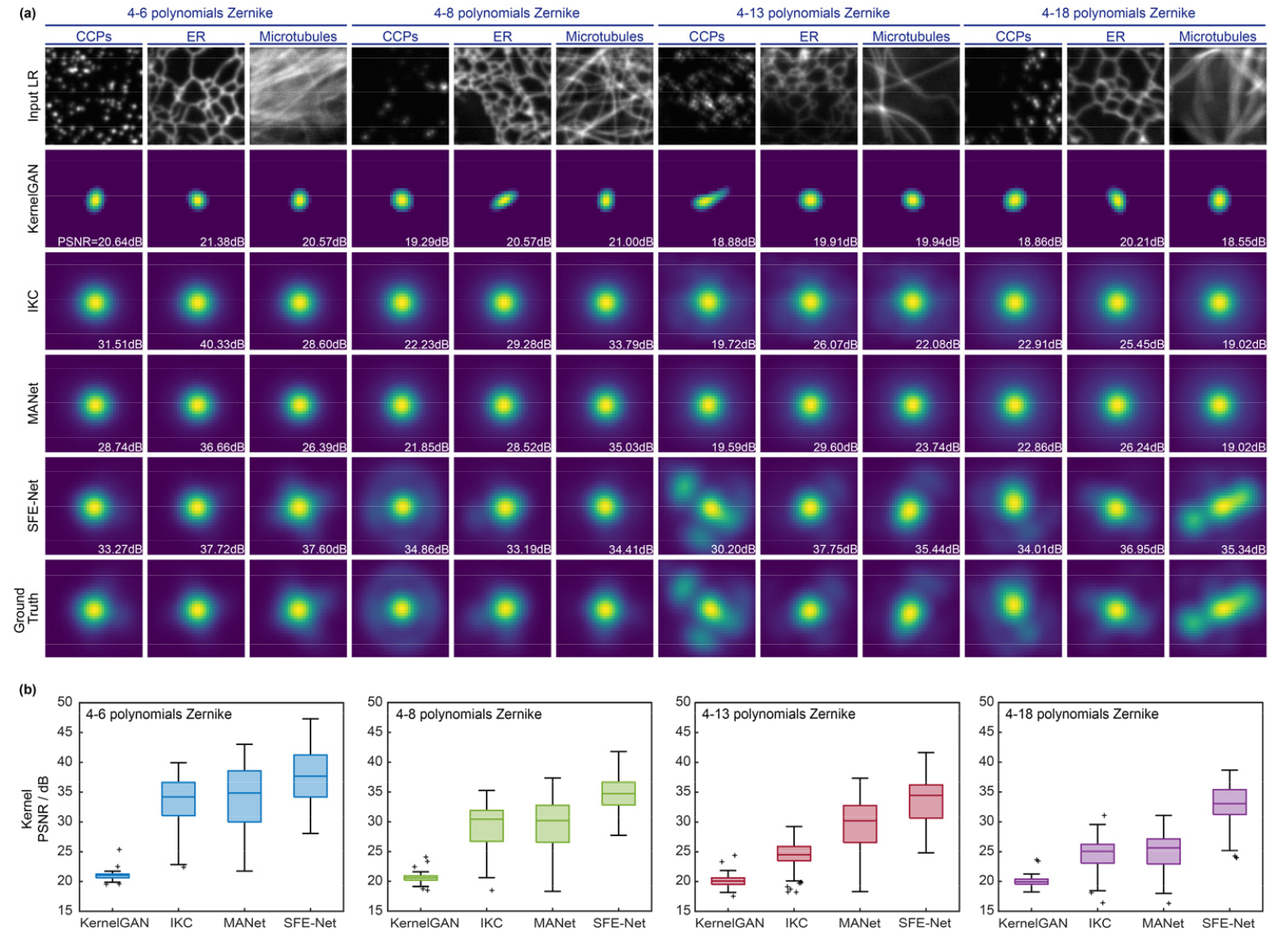
Optical Aberration Estimation via SFE-Net. (a) Representative aberrated PSFs were estimated by KernelGAN, IKC, MANet, and SFE-Net from WF images of CCPs, ER, and MTs. Four groups of datasets with escalating complexity of aberration were generated, corresponding to Zernike polynomials of orders 4-6, 4-8, 4-13, and 4-18. The top and bottom rows show the input WF images and GT PSF images for reference. (b) Statistical comparisons (n=120) of KernelGAN, IKC, MANet and SFE-Net (our proposed method) were conducted in terms of Peak Signal-Noise-Ratio (PSNR) on different training and testing datasets. Center line, medians; limits, 75% and 25%; whiskers, the larger value between the largest data point and the 75th percentiles plus 1.5× the interquartile range (IQR), and the smaller value between the smallest data point and the 25th percentiles minus 1.5× the IQR; outliers, data points larger than the upper whisker or smaller than the lower whisker. The same notations for box plots are used in Fig. 6e and 7b.

To further enhance the feature extraction and representation capability of neural network models in task of PSF estimation, we devised a novel network named Space-Frequency Encoding Network (SFE-Net). SFE-Net leverages both the spatial features and frequential characteristics of the WF image to estimate the aberrated PSF with high accuracy. As shown in Fig. 4, the SFE-Net is trained in a supervised manner to directly map biological WF images to their corresponding aberrated PSFs. Interestingly, before adopting this supervised training scheme, we have gone through a series of physical model-based PSF estimation approaches, such as using untrained network [19], or modified flow-based kernel priors [20]. However, we found that although straightforward, the data-driven supervised mapping strategy with SFE-Net remarkably outperformed other conceptually more complex ideas.

As is shown in Fig. 4a, KernelGAN, IKC, and MANet fail to extract the optical aberration from WF images, even when the aberrated PSF is relatively simple, i.e., generated with 4-6 Zernike polynomials. In contrast, our SFE-Net accurately generates complex aberrated PSF constituted up to 18 orders of Zernike polynomials, with an average peak signal-to-noise ratio (PSNR) higher than 30dB (Fig. 4b). Furthermore, we performed an ablation study on the frequential branch of SFE-Net to validate the gain of incorporating frequential information in the feature extraction process of the PSF estimation network. Specifically, we trained three versions of the SFE-Net on the same training dataset: a standard SFE-Net, a modified version without the frequential branch, and another modified version without the Fast Fourier Transform (FFT) Layer in the frequential branch. The training loss and validation PSNR curves for these three models shown in Fig. 5 demonstrate that the inclusion of the frequential branch, especially the FFT Layer, effectively accelerates network convergence and contributes to an improved PSF estimation performance by 3.2 dB in PSNR.

**Fig. 5.**
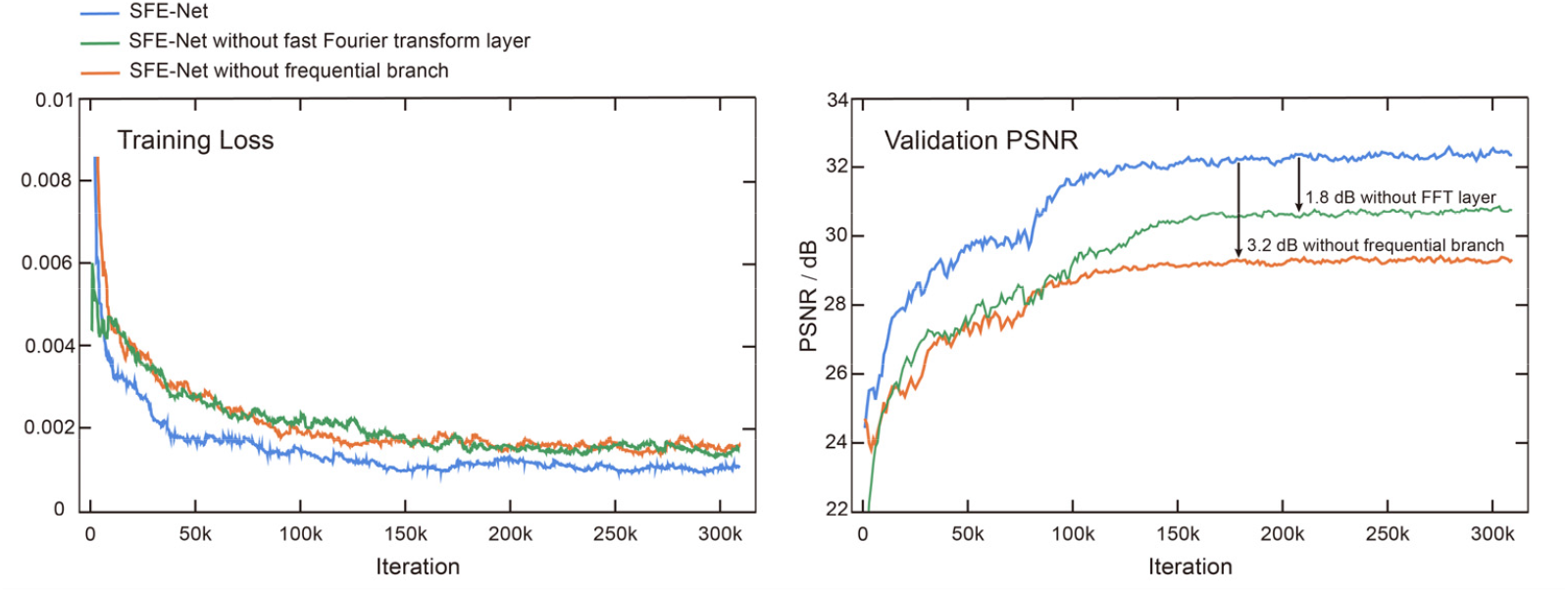
Progression of training loss and validation PSNR of network model with/without frequential branch during training process

### B. Blind Deconvolution with Accurate PSF Estimation

Integrated with deconvolution algorithms, the accurate PSF estimation through SFE-Net provides a straightforward yet efficient solution for numerically compensating optical aberrations and improving spatial resolution in an unsupervised manner. To systematically evaluate the impact of the PSF on deconvolution algorithms during the processing of images with optical aberrations, we processed WF images of CCPs, ER, and MTs with Richardson-Lucy (RL) deconvolution [21, 22] using an ideal Gaussian PSF with theoretically accurate Full Width at Half Maximum (FWHM), as well as PSFs estimated by KernelGAN, IKC, MANet, and SFE-Net (Fig. 6a-c). Our findings demonstrate that RL deconvolution with accurate PSF, i.e., GT PSFs and PSFs estimated by SFE-Net, substantially removes the artifacts induced by aberrations, such as an anomalous distortion in CCP images and ringing artifacts in ER and MT images. Moreover, it enhances both the resolution and contrast for all biological structures. Conversely, when provided with an incorrect PSF estimated by other methods, the RL deconvolution algorithm fails to remove the aberration-induced artifacts and may even generate anamorphic structures (indicated by red arrows in Fig. 6a-c).

**Fig. 6.**
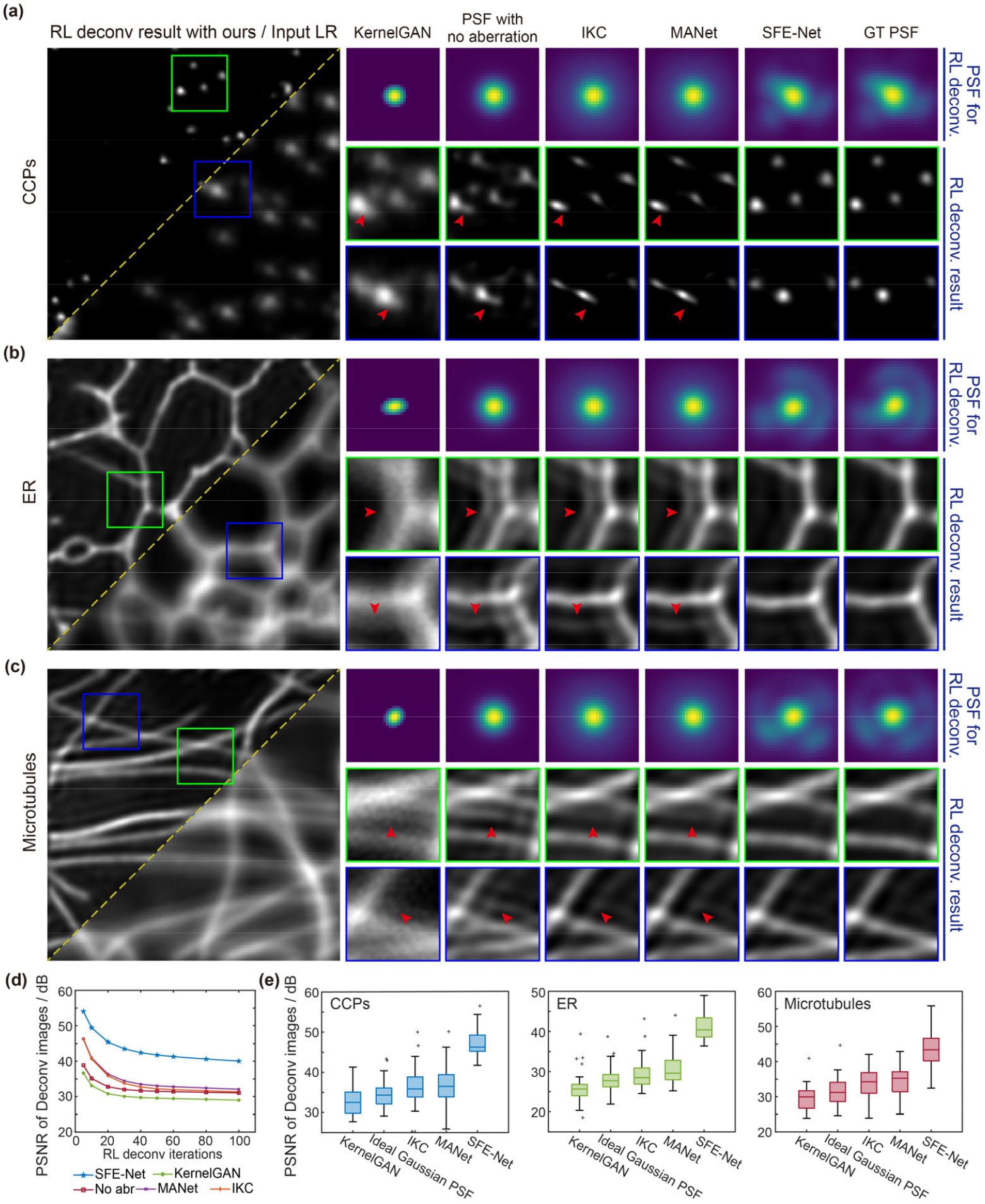
Blind Deconvolution with Estimated PSF. (a-c) Representative deconvolved images of CCPs (a), ER (b), and MTs (c) processed with RL deconvolution algorithm using ideal Gaussian PSF and PSF estimated by KernelGAN, IKC, MANet, and SFE-Net. The aberrated WF images (bottom right in the first column), deconvolved images (top left in the first column) and GT PSF images are shown. (d) PSNR curves calculated between RL deconvolved images using GT PSF and estimated PSFs, with the deconvolution iteration ranging from 5 to 100 (n=120). (e) Statistical comparisons of PSNR for testing datasets of CCPs (left), ER (middle), and MTs (right), respectively (n=30). Scale bar, 1 μm (a-c), 0.5 μm (zoom-in regions of a-c).

We measured the PSNR between deconvolved images obtained using GT PSF and those obtained using estimated PSF, with deconvolution iterations ranging from 5 to 100, across various biological structures. Our results demonstrate that the deconvolved images produced using the PSF estimated by SFE-Net consistently exhibit a significantly higher PSNR compared to those generated using PSFs estimated by other existing methods. This holds true regardless of the number of deconvolution iterations (Fig. 6d) or the specific biological structures being analyzed (Fig. 6e).

### C. Aberration-aware Image Super-resolution

SISR networks have been developed to instantly enhance the resolution of biological images in an end-to-end manner, irrespective of the image formation model [9]. Recent studies have shown that incorporating physical prior knowledges, such as PSF, can improve the performance of super-resolution network. Given the remarkable ability of the proposed SFE-Net to recognize the PSF from low resolution images, we reasoned that incorporating prior knowledge of PSF and optical aberrations could benefit the performance of SISR. To validate this hypothesis, we incorporating the spatial feature transform (SFT) layer [12] with our previously proposed DFCAN model [9] to devise the SFT-DFCAN, which leverages both the aberrated PSF information and Fourier channel attention mechanism to enhance the performance of image super-resolution. Particularly, in SFT-DFCAN, the feature maps are affinely transformed through scaling and shifting operations conditioned by the estimated PSF and aberration, thereby enabling adaptive encoding of the PSF information into the neural networks.Next, we validated the performance of SFT-DFCAN using dataset generated following the steps outlined in Section 2A. We trained a SFT-DFCAN model using pairs of low- and high-resolution image along with their corresponding GT PSFs, and a DFCAN model with low- and high-resolution image pairs for comparison. Fig. 7a displays representative SR images reconstructed using the DFCAN and SFT-DFCAN model with PSFs generated by SFE-Net and other PSF estimation methods. These results show that while a well-trained DFCAN model can partially remove the optical aberration and reconstruct high frequency information, it struggles to capture the fine structure of biological specimens, often resulting in the generation of hallucinated structures (indicated by the red arrow in the sixth column of Fig. 7a). In contrast, the SFT-DFCAN model, benefiting from the prior knowledge of PSF and estimated aberration, has the theoretical capability to recover biological structures with higher fidelity. Both of the qualitative and quantitative comparisons (Fig. 7b) between DFCAN and SFE-Net-guided SFT-DFCAN indicate that the incorporation of PSF and aberration information rationalizes the training and inference process of DFCAN models and provides substantial improvements in output fidelity and resolution.

**Fig. 7.**
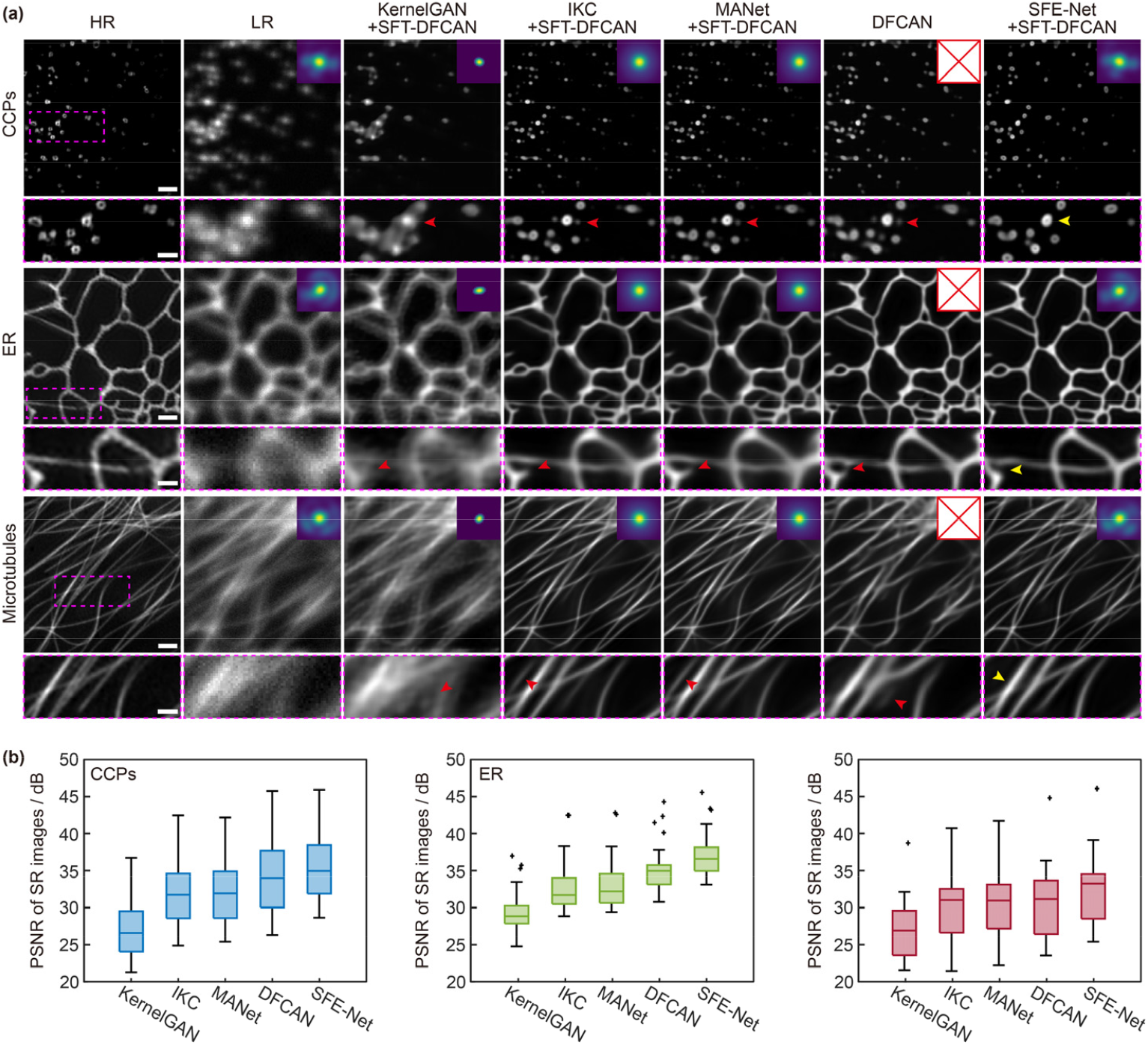
Aberration-aware Image Super-resolution Reconstruction with Estimated PSF. (a) Representative SR images reconstructed by DFCAN and SFT-DFCAN with PSFs obtained from KernelGAN, IKC, MANet and SFE-Net. Low-resolution images and high-resolution GT images are provided for reference. The corresponding estimated PSF images are presented in the top right corner of each reconstructed SR image. Scale bar, 1 μm, and 0.5 μm for zoom-in regions. (b) Statistical comparison of the PSNR values for the output SR images produced by DFCAN and SFT-DFCAN with PSFs estimated by KernelGAN, IKC, MANet, and SFE-Net (n=30).

On the other hand, existing PSF estimation methods such as KernelGAN, IKC, and MANet tend to produce PSF estimates that deviate significantly from the actual one, thereby misleading the high-frequency feature extraction and reconstruction in SFT-DFCAN models. Nevertheless, when facilitated with the aberrated PSF estimated by SFE-Net, the SFT-DFCAN successfully recovers the fine structures of CCPs, ER, and MTs, exhibiting high consistency with the high-resolution GT images (Fig. 7a). Additionally, the statistical comparison of the output SR images generated by different methods (Fig. 7b) demonstrates that SFE-Net-based SFT-DFCAN outperforms other PSF estimation methods-based SFT-DFCAN by a significant margin, enabling high-quality single-image SR reconstruction from aberrated WF images.

### D. Digital Adaptive optics and Super-resolution for Live-cell Imaging

For a well-established optical imaging system, the most common aberrations during live-cell imaging experiments are defocus and spherical aberrations. These aberrations are typically caused by the drifting of the focusing plane, axial movement of the samples, and the misalignment of refractive index between the samples and the cover slip. Additionally, there may be subtle changes or misalignment in the imaging system over its service life, which are often unnoticed and cannot always be corrected in time. To address these inherent optical aberrations in an offline manner, we employed the well-trained SFE-Net and SFT-DFCAN model to perform digital adaptive optics and super-resolution reconstruction for time-lapse experimental WF images.

In our experimental setup, we initially captured 100 consecutive frames of a live SUM159 cell expressing clathrin-EGFP using our home-build multi-modality SIM system. The time interval between each image was set as 0.5 seconds. To simulate focus drifting and axial movement, we deliberately introduced a random disturbance along the z-axis of the motorized sample stage. As depicted in the upper row of Fig. 8a, the WF images of CCPs exhibited varying degrees of defocus aberration at different timepoints. To address this issue, we utilized the proposed SFE-Net to estimate the PSF with varying FWHM for each frame. The estimated PSF was then input into the SFT-DFCAN model, enabling the reconstruction of SR images based on real-time aberrated PSF information. Consequently, the aberrations were effectively removed, leading to the successful resolving of the hollow structure of CCPs (bottom row in Fig. 8a).

**Fig. 8.**
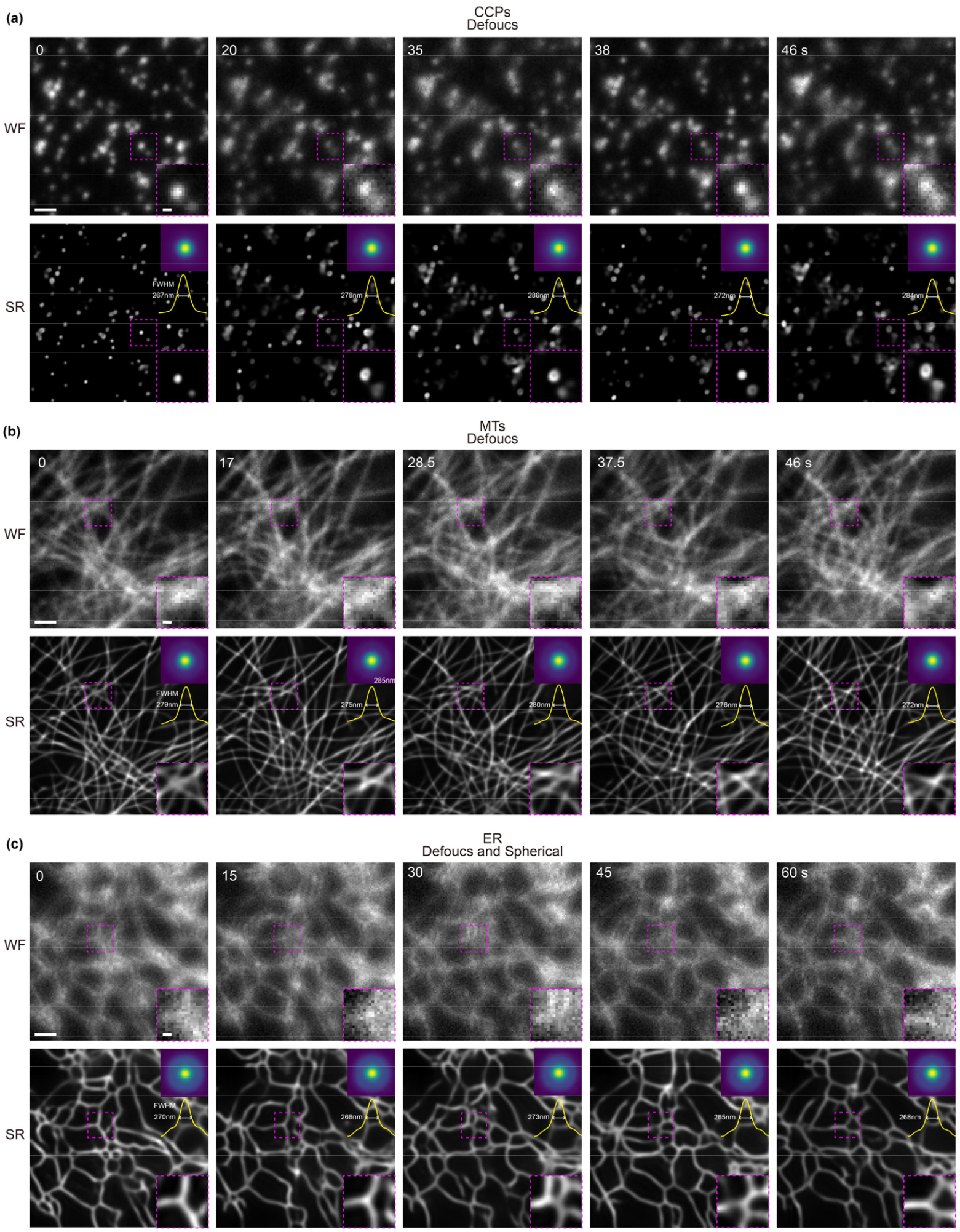
Digital Adaptive optics and Super-resolution for Live-cell Imaging. Time-lapse WF images, estimated PSFs by SFE-Net, and corresponding SR images generated by SFE-Net-facilitated SFT-DFCAN of CCPs (a), MTs (b), and ER (c). During the imaging procedure, the defocus aberration is manually added on CCP (a) and MT (b) data, while a combination of defocus and spherical aberrations are applied on ER images (c). The PSFs estimated by SFE-Net, along with their corresponding profiles and FWHM values, were displayed in the top right corner of SR images. Scale bar, 1 μm (a-c), and 0.2 μm (zoom-in regions of a-c).

Subsequently, we proceeded to image a live COS7 cell expressing Ensconsin-mEmerald and intentionally induced a relatively steady degree of defocus aberration to simulate the misoperation by the operator. Despite severe blurring in the WF images, the SFT-DFCAN equipped with SFE-Net was capable of clearly recovering the densely interlaced microtubules (Fig. 8b). Finally, we imaged another live COS7 cell labelled with calnexin-mEmerald for 100 timepoints, with a time interval of 1 second. Prior to imaging, we deliberately axially offset the sample stage and adjusted the correction collar of the objective to introduce both defocus and spherical aberrations manually. As anticipated, the SFE-Net reliably estimated the time-varying mixed defocus and spherical aberrations. This estimation facilitated the downstream SFT-DFCAN model in resolving the reticular structure of ER with high resolution and contrast (Fig. 8c). These results illustrate that the proposed SFE-Net has the capability to recognize spatiotemporally varying optical aberrations without any additional hardware, except for the aberrated image itself. This capability enables digital aberration compensation and aberration-aware super-resolution reconstruction for time-lapse live-cell imaging.

## 4. DISCUSSION

In this paper, we introduced the SFE-Net, a novel method capable of accurately estimating aberrated PSF directly from WF images. One key advantage of SFE-Net over conventional direct wavefront sensing methods is its ability to provide real-time aberration estimation without requiring any additional optical hardware. Additionally, unlike existing indirect wavefront sensing methods that involves time-consuming iterative acquisition and optimization procedures, SFE-Net can estimate aberrations from a single frame at a timescale of ∼30 millisecond. This makes it suitable for imaging long-term bioprocesses where optical aberrations vary over time and need to be measured and corrected promptly. By utilizing the PSF generated through SFE-Net, we can effectively address various aberrations and improve spatial resolution in biological images using both unsupervised RL deconvolution algorithm and the supervised SFT-DFCAN model. Notably, our experiments revealed that by incorporating prior knowledges of aberrated PSF, the SFT-DFCAN model substantially surpassed its backbone model, DFCAN without embedding PSF information. Finally, we demonstrated the practical applications of SFE-Net and the facilitated SFT-DFCAN model in digitally correcting optical aberrations and achieving instant image super-resolution in time-lapse live-cell imaging experiments.

More potential applications and extension of SFE-Net are anticipated. First, SFE-Net was trained to estimate aberrated PSF generated with up to 18 orders of Zernike polynomials. However, including higher orders of Zernike polynomials could noticeably degrade the performance of SFE-Net. Upgrading the backbone network architecture of SFE-Net to state-of-the-art models such as Swin-Transformer [23] may expand the application scope. Second, in this paper we primarily conducted principal verification of SFE-Net with semi-simulated dataset. However, when applying SFE-Net models trained with simulated data to experimental images, there is inevitably a degradation in performance due to the domain shift problem. More ideally, the training dataset of SFE-Net should be acquired via an imaging system with wavefront shaping capability, and trained with aberrated images experimentally acquired and corresponding ground-truth aberration applied on the wavefront shaping elements. Third, although we mainly demonstrated the offline digital AO functionality of SFE-Net, it can also be used in various hardware-based AO systems for multiple modalities of microscopes or telescopes. In particular, owing to the temporal sensitivity and image patch-based estimation scheme of SFE-Net, it can be applied to measure and correct the drastically varying aberrations both spatially and temporally in telescope technologies. We hope that our methods will inspire further developments of next-generation adaptive optics and super-resolution microscopy.

## Acknowledgments

The authors thank T. Kirchhausen for the donor plasmids used for genome editing and help in generating the genome-edited cell lines.

## Data availability

The python codes of SFE-Net and the MATLAB codes for generating training and testing dataset with BioSR are publicly accessible on Github (https://github.com/HypnosRin/SFE-Net). Data underlying the results presented in this paper are available upon reasonable requests.

